# Ecological outcomes of hybridization vary extensively in *Catostomus* fishes

**DOI:** 10.1101/2021.01.20.427472

**Authors:** Elizabeth G. Mandeville, Robert O. Hall, C. Alex Buerkle

## Abstract

Hybridization has been studied extensively to learn about speciation and mechanisms of reproductive isolation, but increasingly we recognize that hybridization outcomes vary geographically and can depend on the environment. At the same time, hybridization can reshape biotic interactions in an ecosystem, leading to ecological shifts where hybridization occurs. Identifying how hybrid individuals function ecologically would link evolutionary outcomes of hybridization to ecological consequences, but relatively few studies have focused on ecological traits of hybrids. We described the feeding ecology of several *Catostomus* fish species and their hybrids by using stable isotopes (*δ*^13^C and *δ*^15^N) as a proxy for diet and habitat use, and compared two native species, an introduced species, and three interspecific hybrid crosses. We replicated this comparison spatially, including hybrids and parental species from seven rivers in the Upper Colorado River basin where parental species co-occur and the opportunity for hybridization exists. Although relationships between native species in isotopic space varied, individual native species did not fully overlap in isotopic space in any river sampled, suggesting little overlap of resource use between these historically sympatric species. The introduced species overlapped with one or both native species in every river, suggesting similar resource use and potential for competition. Hybrids occupied intermediate, matching, or more extreme (transgressive) isotopic space relative to parental species, and were isotopically variable within and among rivers. We suggest that ecological outcomes of hybridization, like genomic outcomes of hybridization, are likely to vary across locations where a pair of species hybridizes. This variation implies that hybridization might have large unpredictable, idiosyncratic ecological effects on fish assemblages where hybrids occur. Although we found little evidence that hybrids are at a disadvantage ecologically—there were no significant declines in body condition relative to parental species—it is nevertheless possible that abiotic or biotic attributes of a river might constrain the range of interspecific hybrids that are successful, thus contributing to variation in hybridization outcomes across rivers.

## Introduction

Hybridization is a critical feature of many speciation processes and serves as a test of reproductive isolation between diverged lineages in secondary contact (Barton and Hewitt 1989, Harrison 1990). Studies of hybridization are motivated in part by a desire for generality in our understanding of how biodiversity arises and is maintained. While the increasing accessibility of large genomic datasets for wild populations has led to a newfound ability to study replicate outcomes of hybridization in the wild (Narum et al. 2013, McFarlane and Pemberton 2019), better information in some cases leads to a more nuanced and complicated understanding of evolution, and generates additional questions (Payseur and Rieseberg 2016). Studies of hybridization in multiple locations have uncovered substantial variation in genomic outcomes of hybridization when a pair of species comes into contact repeatedly (Nolte et al. 2009, Lepais and Gerber 2011, Haselhorst and Buerkle 2013, Lagache et al. 2013, Mandeville et al. 2017), and it seems increasingly likely that ecological context, historical contingency, and stochasticity influence to what extent species remain reproductively isolated from their close relatives. Variable outcomes across replicate instances of hybridization between the same pair of species suggest not only variation within a species in mechanisms of reproductive isolation (Cutter 2012, Mandeville et al. 2017), but also variable porosity of boundaries to gene flow between species. Identifying the degree of ecological dependency and contingency in hybridization outcomes is therefore essential to understanding speciation and the persistence of biodiversity.

Mechanisms generating variation in hybridization outcomes remain unidentified in most hybridizing species pairs (Gompert et al. 2017). Ecological interactions and environmental conditions likely contribute significantly, and indeed, empirical examples of environmental dependence are known (e.g. Taylor and Donald McPhail 2000, Young et al. 2016, Muhlfeld et al. 2017, Mandeville et al. 2019). However, our understanding of the ecological context and consequences of hybridization is still largely incomplete, which is unfortunate, because ecological context can influence outcomes of hybridization both by affecting relative fitness of hybrid and parental individuals (e.g. Arnold et al. 2012) and by limiting or increasing the opportunities for hybridization (e.g. Lepais and Gerber 2011). Fitness and ecological success of hybrids might also vary across independent instances of hybridization, with some phenotypes and genomic compositions of hybrids being favored in a subset of locations, and alternative traits and genotypes being favored in others. Differential ecological success of hybrids across replicate hybrid zones could selectively filter which genotypes persist (Lindtke et al. 2014), and might contribute to variation in hybridization outcomes across locations. However, few empirical studies have described both variation in hybridization outcomes and the ecological conditions that have either resulted from or shaped these outcomes.

In fish, ecological success of interspecific hybrids can depend on environmental conditions (e.g. in sticklebacks; Hatfield and Schluter 1999). The presence of hybrids can alter species interactions so growth, survival rate, or reproductive success of parental species is changed where hybridization occurs (as in trout; Rosenfield et al. 2004, Seiler and Keeley 2009). Phenotypes related to foraging ability can be especially important in determining outcomes of interactions between traits of hybrids and the environment. Experimental work in fish has shown that hybrids are sometimes less successful than parental species when they have intermediate mouth morphology, because of an ecological mismatch between feeding morphology and available resources, leading to worse feeding performance (e.g. sunfish; McGee et al. 2015) or reduced growth (e.g. sticklebacks; Arnegard et al. 2014). However, phenotypes of hybrids are hard to predict, and are not always intermediate between parental species (Rieseberg et al. 1999*b*, Stelkens et al. 2009). Instead, phenotypes of hybrid individuals might be similar to one of the parental species, or might differ from both parental species (e.g., Stelkens et al. 2009, Thompson et al. 2019).

Our goal for this study was to compare ecological interactions between multiple *Catostomus* fish species and their hybrid progeny across several rivers in Colorado and Wyoming. The focal *Catostomus* species are part of a clade with extensive and highly variable interspecific hybridization between multiple species pairs (Hubbs et al. 1943, McDonald et al. 2008, Mandeville et al. 2015, 2017). We used stable isotopes of carbon and nitrogen as a proxy for diet and spatial resource use, and assessed patterns of ecological overlap first between parental species, and then between parental species and hybrid individuals. We compared patterns of ecological overlap across multiple geographic locations locations where fish were sampled, and analyzed the relationship between these ecological overlaps and genomic outcomes of hybridization in these rivers.

Genomic outcomes of hybridization in *Catostomus* can vary substantially over the geographic range in which a pair of species co-occurs (Mandeville et al. 2015, 2017). While some hybridization between *Catostomus* species is precipitated by human-caused species introductions, most notably of *C. commersonii* (white sucker), this group of fishes likely has a long and convoluted history of hybridization (Hubbs et al. 1943, Hubbs 1955, Bangs et al. 2018), including a hypothesized allopolyploid origin (Ferris 1984). Contemporary hybridization between six geographically overlapping *Catostomus* species is extensive; several species have hybridized with more than one congener, resulting in at least 12 different hybrid crosses (Mandeville et al. 2017). In some locations, few crosses were present despite sympatry of multiple parental species; in other locations, many different crosses involving the sympatric parental species occurred. Backcrossed hybrids and later-generation recombinant hybrids were common in some locations, while other locations contained only first generation hybrids (Mandeville et al. 2015, 2017). Mechanisms driving this variation in hybridization outcomes remain unidentified. Environmental variation that affects opportunity for hybridization or intrinsic genetic variation in reproductive isolation could explain the variable hybridization outcomes that we observed in natural populations (Mandeville et al. 2017). Another possibility, which we explore in this study, is that geographic variation in the ecological success and survival of hybrids might cause variation in hybridization outcomes.

One focal species, bluehead sucker (*Catostomus discobolus*), eats predominantly algae that grow in fast-moving, shallow portions of rivers (i.e., riffles), while the other two widespread parental species, flannelmouth and white suckers (*C. latipinnis* and *C. commersonii*), eat more aquatic invertebrates and more often occur in deep, slow pools (Tronstad and Estes-Zumpf 2011, Cross et al. 2013, Walsworth et al. 2013). *Catostomus* species vary substantially in mouth morphology. *C. discobolus* and *C. platyrhynchus* have a distinctive “scraping ridge” that facilitates scraping algal biofilms from surfaces (Baxter et al. 1995). In contrast, the other species have no scraping ridge, but have fleshy lips in varying shapes and with different textures. Differences in diet and foraging location among species and hybrid crosses can be captured using stable isotopes. Stable isotopes of nitrogen (*δ*^15^N) serve as a proxy for trophic position, and would therefore capture the difference between consuming algae and invertebrates; we expect higher values of *δ*^15^N for individuals consuming more invertebrates, because they are feeding on a higher trophic level (Post 2002). Stable isotopes of carbon (*δ*^13^C) capture differences in carbon composition in diet, which can derive from intrinsic characteristics of dietary items (e.g. C3 versus C4 plants), but in rivers might also depend on patterns of CO_2_ exchange between water and air, source of CO_2_, and flow velocity over biofilms (Finlay 2001, Newsome et al. 2007). Carbon stable isotopic ratios might therefore also differ depending on how much individuals feed in riffles or pools in a river. Separation of isotopic niches, as detected by stable isotope analysis, might be a key variable in persistence of closely related species, as in other fish species (e.g. sauger and walleye; Butt et al. 2017). We used stable isotope data to identify ecological overlaps between genetically-defined groups (Layman et al. 2007, Newsome et al. 2007). We compared ecological success of hybrids and parental species using body condition as a proxy for fitness and described the success of parental and hybrid fish in the context of their isotopic niche and overlap with other species or hybrids.

## Methods

We combined stable isotope and genetic analyses of 506 individual fish to identify how ancestry of *Catostomus* suckers is related to ecological niche. Sampling spanned seven different locations in the Upper Colorado River basin in Wyoming and Colorado (Table 1). Fisheries biologists from state agencies collected fin tissue from each individual fish. Fin tissue was used for both genetic analyses (Mandeville et al. 2015, 2017) and stable isotope analyses (this study). Locations for isotope analysis were chosen to include rivers with a range of hybridization outcomes, based on the results of Mandeville et al. (2017).

**Table 1:** Fish were sampled from seven rivers in the Upper Colorado River basin. Individuals sampled included parental species and hybrids, abbreviated in the table below. (B = blue-head, F = flannelmouth, W = white. Hybrids are represented with a *×* connecting parental species, i.e. flannelmouth*×*white = F*×*W).

| Location | Drainage | Individuals | Species | Hybrids (> 3 ind) |
| --- | --- | --- | --- | --- |
| Big Sandy River (WY) | Green River | 61 | B, F, W | — |
| Little Sandy Creek (WY) | Green River | 80 | B, F, W | F×W |
| Escalante Creek (CO) | Gunnison | 85 | B, F, W | B×F, B×W, F×W |
| White River (CO) | White | 55 | B, F | B×F |
| Little Snake River (WY) | Yampa | 68 | F, W | B×W, F×W |
| Yampa River (CO) | Yampa | 72 | B, F, W | B×W, F×W |
| Muddy Creek (WY) | Yampa | 58 | B, W | B×W, F×W |

We included individuals from five *Catostomus* fish species, *C. discobolus* (bluehead sucker), *C. latipinnis* (flannelmouth sucker), *C. commersoni* (white sucker), *C. catostomus* (longnose sucker), and *C. platyrhynchus* (mountain sucker). Three focal species—bluehead, flannelmouth, and white suckers—are geographically widespread in the Upper Colorado River basin and were present at most sampling sites, whereas longnose and mountain suckers were sampled in only one location each, but were included because they are part of food web interactions where present. Of the five species included in this study, bluehead, flannelmouth, and mountain suckers are native to the Upper Colorado River basin, and white and long-nose suckers were introduced in the Upper Colorado River basin (Baxter et al. 1995). It is unknown exactly when and how white and longnose suckers were introduced, but the introduction probably occurred in the past 100–150 years (Gelwicks et al. 2009, Senecal et al. 2010). Recent work has focused on hybridization between the introduced white sucker and native bluehead and flannelmouth suckers (McDonald et al. 2008, Mandeville et al. 2015), but hybridization also sometimes occurs between the common native *Catostomus* species, bluehead and flannelmouth sucker (Hubbs et al. 1943, Mandeville et al. 2017). Additionally, in some locations mountain and longnose suckers hybridize with the more common species. Hybridization produces spatially heterogeneous outcomes, and different hybrids were sampled in different rivers, with variable amounts of backcrossing to parental species (Mandeville et al. 2015, 2017, Fig. 1).

**Figure 1:**
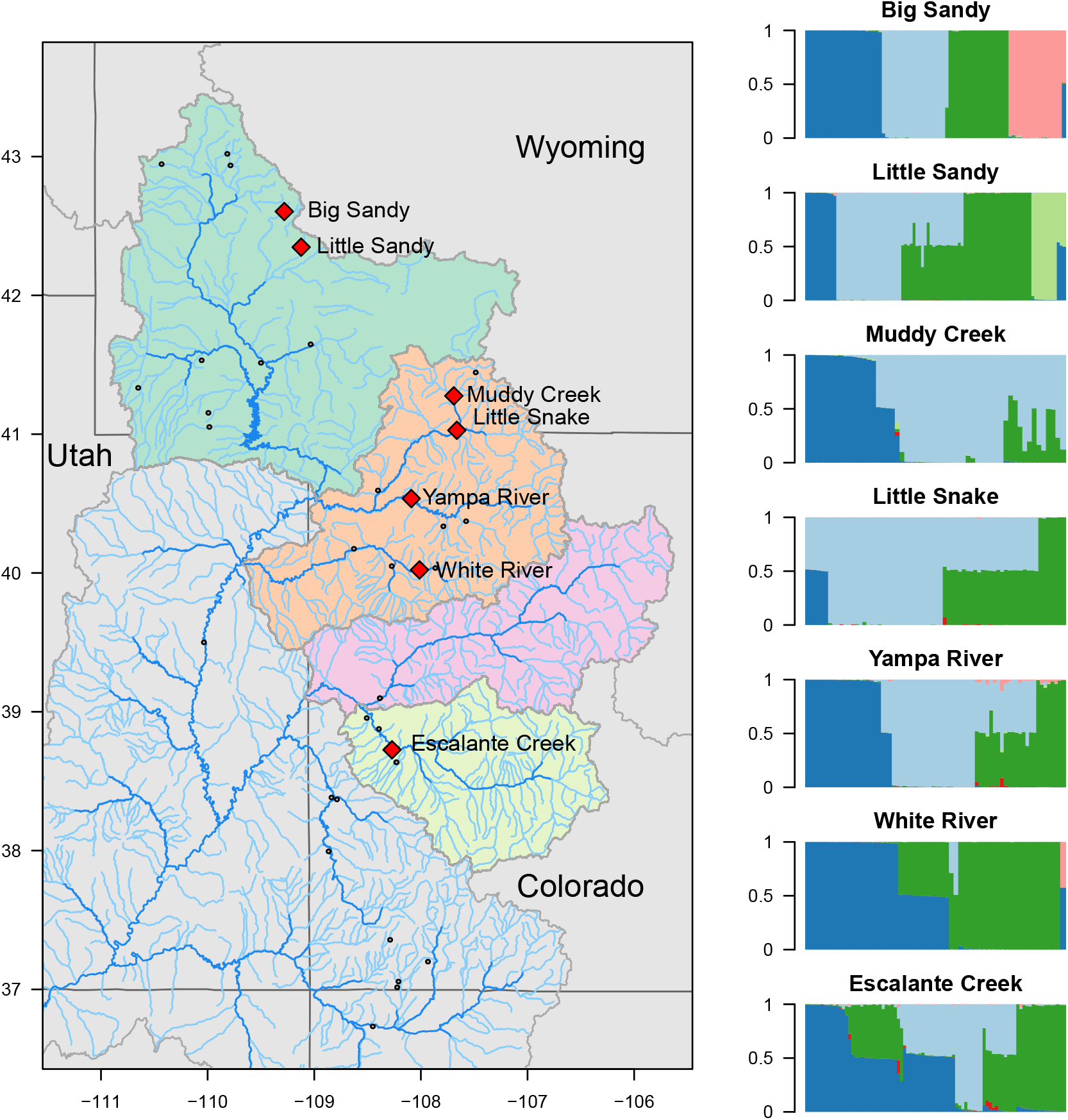
Seven rivers in Wyoming and Colorado and corresponding genetics outcomes of hybridization had indicated that a range of hybrids between *Catostomus* species were present (Mandeville et al. 2017). Gray dots represent all sampling points from Mandeville et al. (2017); the focal rivers for this study are shown with red diamonds. Barplots at right show proportional genetic ancestry of individual fish used for stable isotope analysis (dark blue = bluehead sucker; light blue = white sucker; dark green = flannelmouth sucker; pink= longnose sucker; light green = mountain sucker).

We used results of genomic analyses from Mandeville et al. (2017) to identify ancestry of individuals (Fig. 1). To briefly summarize, genomic analyses of hybridization for the individuals in this study used genotyping-by-sequencing data (Parchman et al. 2012). Specifically, we relied on results from a genetic clustering algorithm, entropy (Gompert et al. 2014, Shastry et al. 2021), based on 11,221 single nucleotide polymorphisms (SNPs) to identify parental species and hybrids genetically. This analysis produced precise estimates of each individual fish’s ancestry, including proportion of ancestry from each parental species in hybrids. Estimates of interspecific ancestry from entropy also revealed extent of hybridization and backcrossing, so we know that a substantial proportion of hybrids in this study were *F*_1_ (first generation) hybrids or *BC*_1_ (backcrosses resulting from mating between an *F*_1_ hybrid and a parental individual). We used both categorical designations of ancestry (species or hybrid cross) and a continuously valued measure of ancestry (proportion of ancestry) as covariates for stable isotopic ratios and body condition of hybrids in this study. More extensive discussion of genomic outcomes of hybridization in *Catostomus* fishes can be found in Mandeville et al. (2017).

To obtain stable isotope data, we prepared samples of fin tissue for analysis of *δ*^13^C and *δ*^15^N by first desiccating them in a drying oven (60 ^°^C) for 2–3 days. Dried fin samples were analyzed the University of Wyoming Stable Isotope Facility (UW-SIF). UW-SIF ground dried fin into a powder, and then transferred an appropriate mass of sample into a tin. We then measured *δ*^13^C and *δ*^15^N ratios using a Finnigan DeltaPlus XP mass spectrometer connected to a Costech 4010 elemental analyzer. UW-SIF normalized values using known laboratory standards (keratin), and performed quality control on all data.

For each species or hybrid cross in each river, as defined by estimates of genetic ancestry (Mandeville et al. 2017), we estimated an ellipse in dual isotopic space to quantify the relative position of *δ*^13^C and *δ*^15^N values and variation in those values within a group (Layman et al. 2007). We used the maximum likelihood `standard.ellipse` function in the `siar` package in `R` to fit the standard ellipses (Parnell et al. 2010, Jackson et al. 2011). This model assumes that bivariate isotopic data (isotopic ratios of *δ*^13^C and *δ*^15^N) for a group of individuals is best fit by a multivariate normal distribution, and uses sample means and a covariance matrix to estimate parameters of a standard ellipse containing approximately 40% of the data (Jackson et al. 2011). To account for potential underestimation of ellipse size at low sample sizes, a correction of (n-1)/(n-2) is applied to produce SEAc, a standard ellipse that is unbiased with respect to sample size. We used the SEAc version of the standard ellipse for all analyses, because simulations suggest that it is less sensitive to sample size variation and small sample size (Jackson et al. 2011), and our sample sizes varied across rivers and species sampled (Table 1). We quantified isotopic similarity of parental species and hybrids within each river using the `overlap` function in `siar` to calculate the overlap of standard ellipses (Jackson et al. 2011). Because rivers likely varied in baseline *δ*^13^C and *δ*^15^N ratios, as well as the food resources available, we calculated overlaps only within a river, not across rivers. We then compared area of isotopic overlap in each pair of species or hybrids across rivers to understand how consistent relative positions of species are in isotopic space.

We also assessed the distribution of individuals in isotopic space using a hierarchical Bayesian model, `isoclust`, written in `JAGS` and implemented through R (Code included in Supplement). This model identifies the best-supported number of distinct clusters in isotopic space for each river, thus quantifying how clustered individuals are in resource use. We sequentially fit models for 1 to 6 clusters in isotopic space, where each individual is assigned categorically to a cluster, and identified the best fit of model to data using a penalized measure of model deviance (pD). This approach complements our more conventional analyses where we quantified resource use by binning individuals a priori by species. By identifying statistically how many clusters are represented in isotope space, we can better quantify distinctiveness and overlap across species and hybrid categories, rather than simply describing clusters qualitatively based on the assumption that individuals with similar ancestry will be ecologically similar (i.e., grouping by ancestry as in Fig. 2).

**Figure 2:**
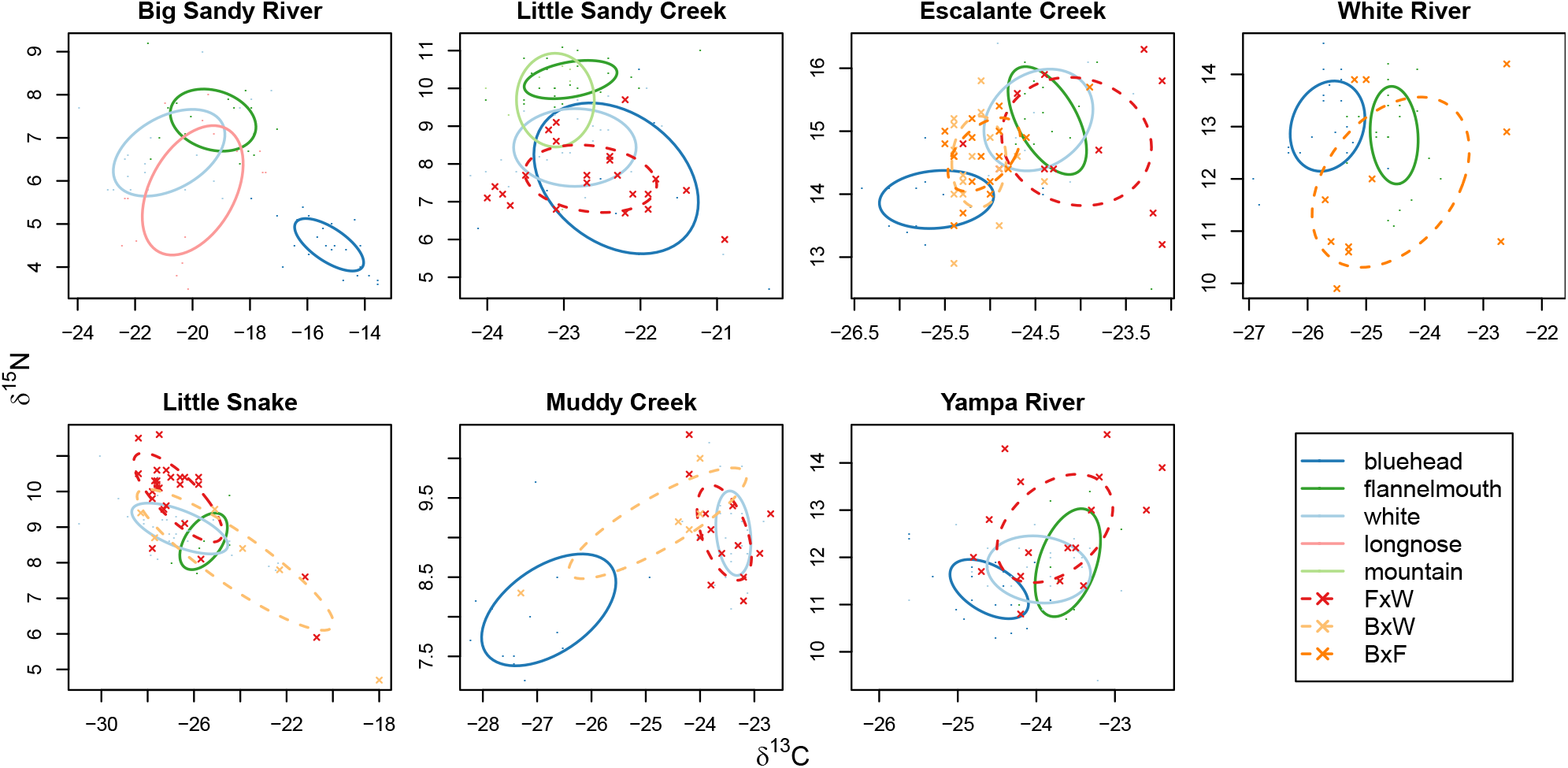
Standard ellipses summarize stable isotopic ratios (*δ*^13^C and *δ*^15^N) by encompassing approximately 40% of individual fish within each of five species and three hybrid crosses. Ellipses were estimated using a maximum likelihood and a bivariate normal distribution, implemented in SIAR (Jackson et al. 2011). Species ellipses are solid lines; hybrid ellipses are dashed lines. Ecological relationships between parental species varied across rivers, but the two native parental species (bluehead and flannelmouth) did not overlap in isotopic space in any sampled locations. Hybrids among these three parental species had matching, intermediate, or transgressive isotopic signatures relative to parental species.

Finally, to characterize relative ecological success of parental species and hybrids, we calculated body condition using length and weight data for 255 individual fish that had these data available (Bolger and Connolly 1989, Jakob et al. 1996). Body condition is a fitness proxy, but because hybrids are likely to vary in fertility independent of their body condition, we interpret body condition as a correlate of ecological success but not necessarily of fitness. We used two methods, Fulton’s condition factor (K; Bolger and Connolly 1989) and relative weight (W_*r*_; Wege and Anderson 1978). We first used Fulton’s condition factor (Anderson and Neumann 1996) because it is simple and requires no species-specific constants, allowing use on hybrids as well as parental species. The equation we used is *K* = (*W/L*^3^) *×* 100, 000. To avoid distortion due to ontogenetic body shape changes, we excluded individuals *<*200 mm in total length. Although body condition can be affected by differences in body shape (e.g., degree of lateral compression), we assumed that the *Catostomus* species in this study are similar enough in shape to compare body condition across species within the genus using Fulton’s condition factor. The only major difference in body shape among these species is that white suckers have a wider caudal peduncle, and bluehead and flannelmouth suckers have a more slender caudal peduncle (Baxter et al. 1995). We also used relative weight (W_*r*_) to compare body condition for flannelmouth (Didenko et al. 2004) and white suckers (Bister et al. 2000), the two species with standard weight equations available (98 individual fish). These equations were developed based on typical length and weight for reference individuals in each species. W_*r*_ is thought to be better than Fulton’s condition factor for comparing across species with different body shapes, but is also limited because not all species in this study could be included. There are not yet published equations for W_*r*_ for bluehead suckers, and more problematically, it is unclear which equation should be used for hybrid individuals or indeed whether any of the species-specific equations are appropriate for hybrid individuals. Hybrids were not included in W_*r*_ analyses, both because there is no developed equation, and because hybrids are likely more phenotypically heterogeneous than individuals from parental species, so applying an equation from a parental species would be inappropriate.

## Results

Flannelmouth suckers and bluehead suckers, which are both native to the rivers we studied, did not overlap in bivariate isotopic space (carbon and nitrogen isotopes) in any of the five sampling locations where both species were sampled (Fig. 2), suggesting minimal overlap of resource use. In two locations (Big Sandy River and Escalante Creek), these two species were substantially separated in dual isotope plots, whereas in the other three locations, standard ellipses for these species were directly adjacent but not overlapping (Fig. 2). Flannelmouth suckers were more enriched in nitrogen than bluehead suckers in four of five rivers where both species were sampled, suggesting that flannelmouth suckers feed on a higher trophic level. However, in one river (Little Sandy Creek), flannelmouth suckers were differentiated from blueheads only in *δ*^15^N, not in *δ*^13^C. In another location (White River), flannelmouth and bluehead suckers were differentiated only in *δ*^13^C, not in *δ*^15^N. In each river in this study, white sucker, a widespread introduced species, overlapped in dual isotope plots with at least one of the native species (Fig. 2). In four locations (Big Sandy River, Escalante Creek, Little Snake River, and Muddy Creek), white suckers overlapped with flannelmouth suckers in isotopic space, and in one location, white suckers overlapped with bluehead suckers (Little Sandy Creek). In one location, white suckers overlapped with both parental species (Yampa River). The variation in overlap between native and introduced species suggests different ecological effects of the white sucker introduction in different locations.

The extent of backcrossing in flannelmouth*×*white sucker hybridization correlated negatively with the overlap in isotopic niche in a river; there was less backcrossing of hybrids to parental species in rivers with more ecological overlap between species, and more backcrossing in rivers with less ecological overlap between flannelmouth and white suckers (Fig. 4B). Relative resource use of hybrids and parental species varied. In one river (Escalante Creek), bluehead*×*white and bluehead*×*flannelmouth hybrids had intermediate isotope values between parental species for both carbon and nitrogen (Fig. 2, Fig. 5), and also slightly overlapped with both parental species. In another river (White River), bluehead*×*flannelmouth hybrids overlapped with parental species in carbon isotopic ratio, but had extremely variable nitrogen ratios that were sometimes much lower than either parental species. Flannelmouth*×*white hybrids overlapped substantially in dual isotope plots with both white and flannelmouth suckers in three out of four rivers in this study where flannelmouth*×*white suckers were sampled, suggesting that some hybrid individuals might use similar food resources to parental species. In two locations (Escalante Creek and the Yampa River), a subset of flannelmouth*×*white hybrid individuals also had transgressive isotopic ratios (Fig. 6), or isotopic ratios that substantially exceeded the range of either parental species and were not intermediate between parental species. In Escalante Creek, these transgressive hybrids were more enriched in carbon than parental species, suggesting that they might use a different carbon source, consistent with foraging in a different location. In the Yampa River, transgressive hybrids were more enriched in nitrogen, suggesting that they might have foraged on a higher trophic level.

For the hierarchical Bayesian clustering analysis, models with 1–3 clusters best fit the isotopic ratio data for individual fish in each river (Fig. 3). However, these clusters of ecologically similar individuals did not correspond to species or hybrid crosses. The number of statistically supported clusters (1–3) was correlated with the number of species (2–5) sampled in each river (*p* = 0.0457; Pearson correlation 0.764), but in each location, fewer clusters were identified in isotopic space than there were species present (Fig. 4A). There was no relationship between number of genetic categories (species plus hybrid crosses) and the number of statistically defined clusters in isotope space. Additionally, with the exception of bluehead suckers in the Big Sandy River and Muddy Creek, clusters defined by our model did not correspond to species (Fig. 3). Within a river, each cluster contained multiple species; similarly, individuals of the same species were typically spread across more than one model-defined cluster.

**Figure 3:**
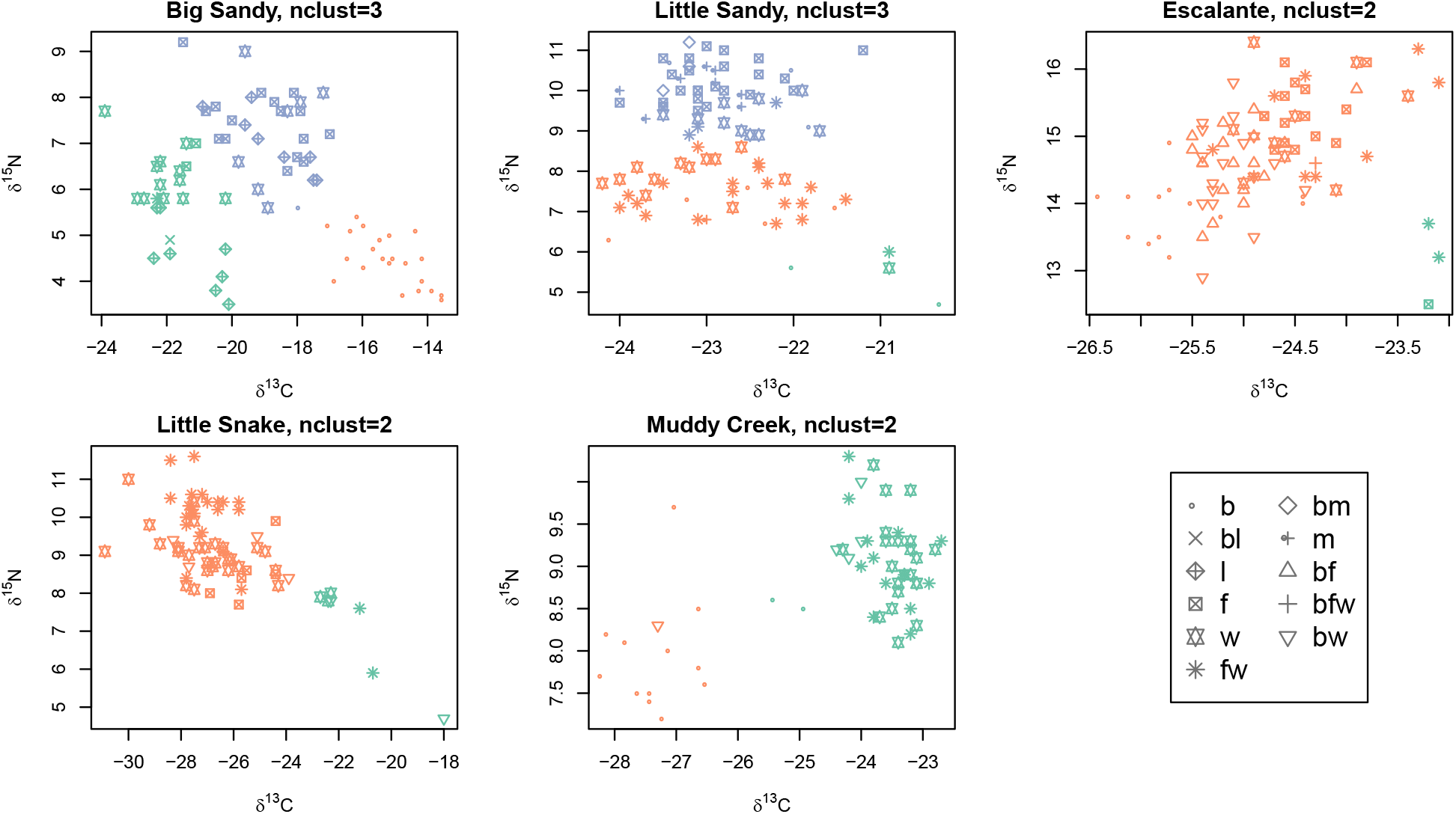
Results of a hierarchical Bayesian clustering algorithm were plotted in dual isotopic space for the five rivers where the optimal number of clusters was greater than 1. Point color corresponds to cluster membership as estimated by our model; point shape corresponds to species or hybrid cross. Estimates of the optimal number of clusters in isotopic space and membership of individuals in those clusters suggest that defined clusters did not correspond closely to species or hybrid categories in most cases.

**Figure 4:**
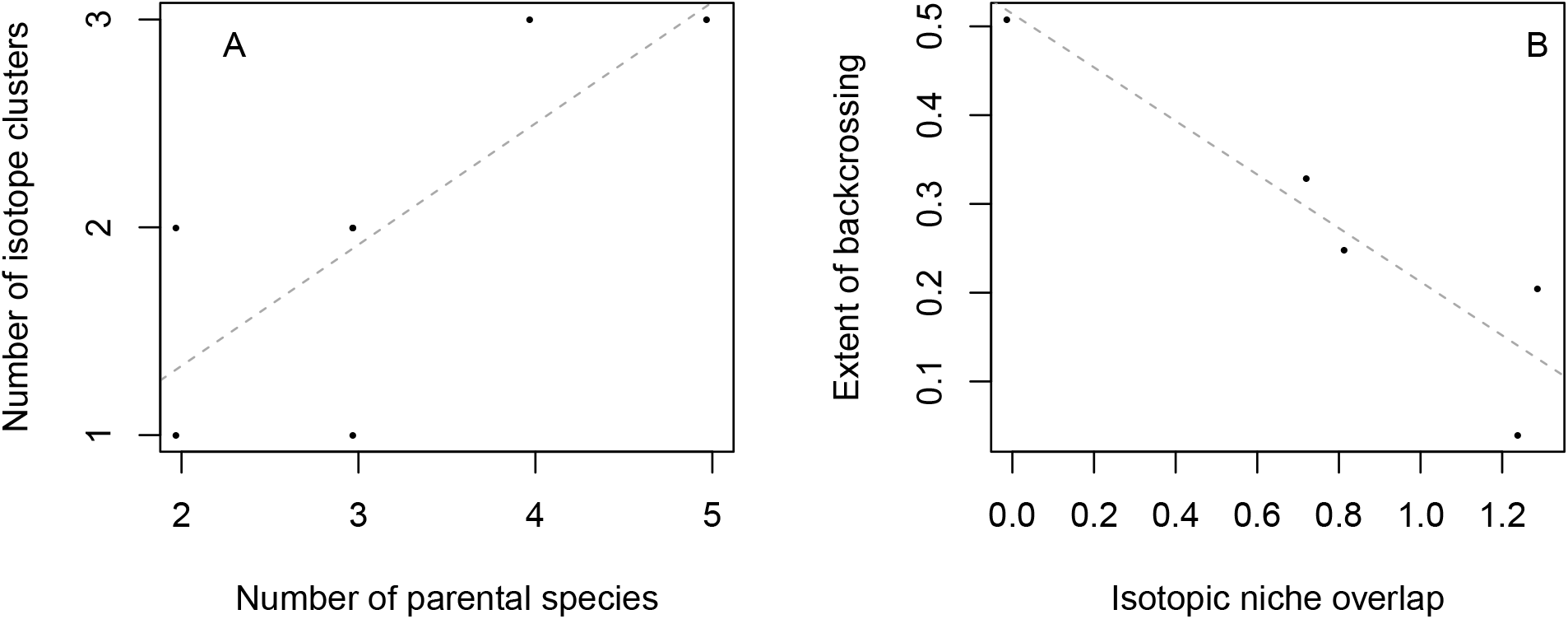
A) Number of clusters of ecologically similar individuals (identified by a hierarchical Bayesian model) correlated with the number of parental species sampled in a river. B) Extent of backcrossing in flannelmouth × white hybrids negatively correlated with the degree of isotopic niche overlap in 6 river locations.

**Figure 5:**
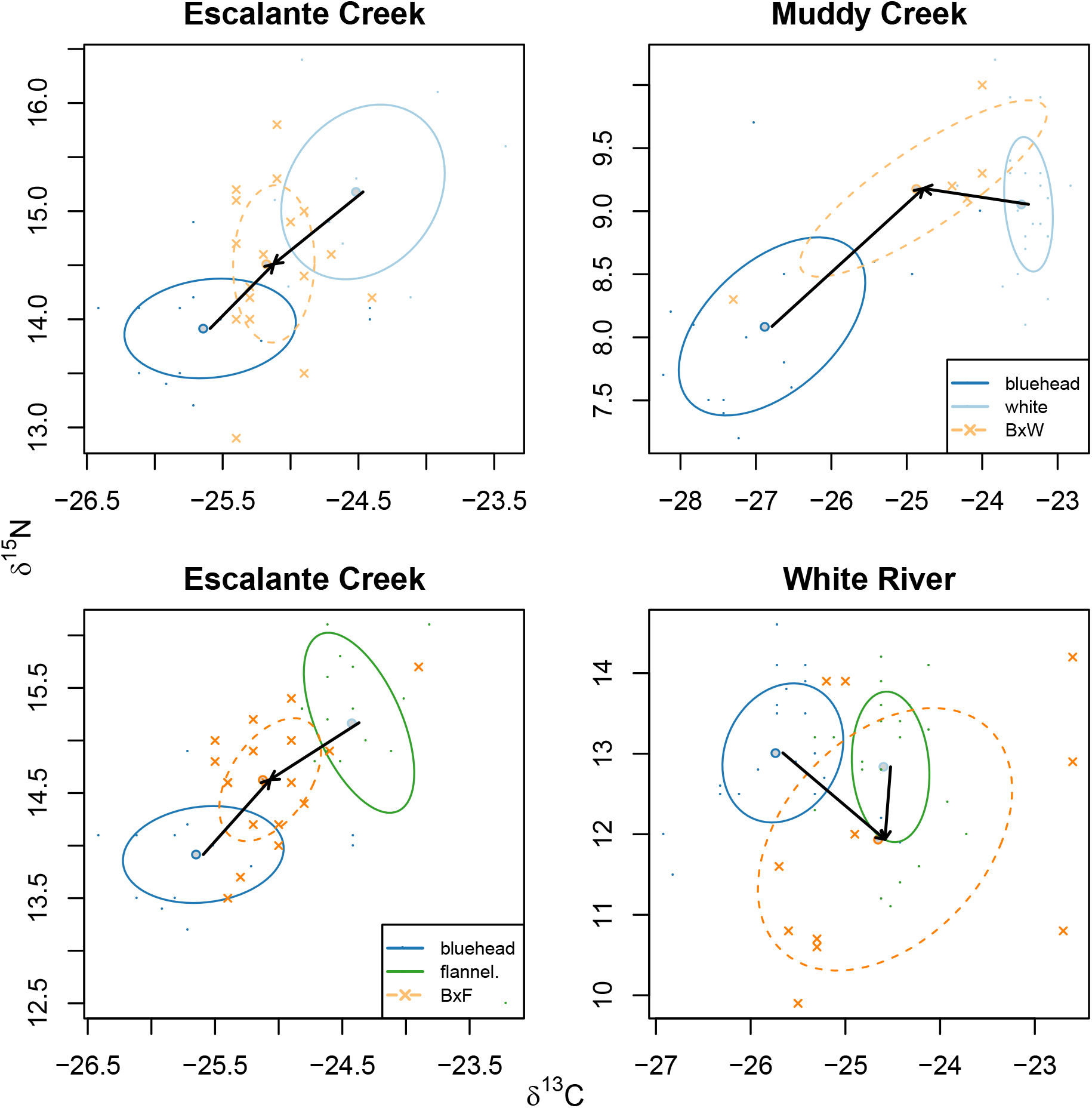
Mean carbon and nitrogen isotopic ratios for bluehead × white sucker hybrids (top panels) were intermediate between values for the two parental species in both carbon and nitrogen, as shown by arrows from the mean for parental species to the mean for hybrid individuals. Mean carbon and nitrogen isotopic ratios for bluehead × flannelmouth sucker hybrids (bottom panels) were intermediate between values for the two parental species Escalante Creek, but transgressive relative to parental species in the White River as shown by arrows from the mean for parental species to the mean for hybrids. Means are shown with gray points; individual fish are denoted by points color-coded by species or hybrid cross.

**Figure 6:**
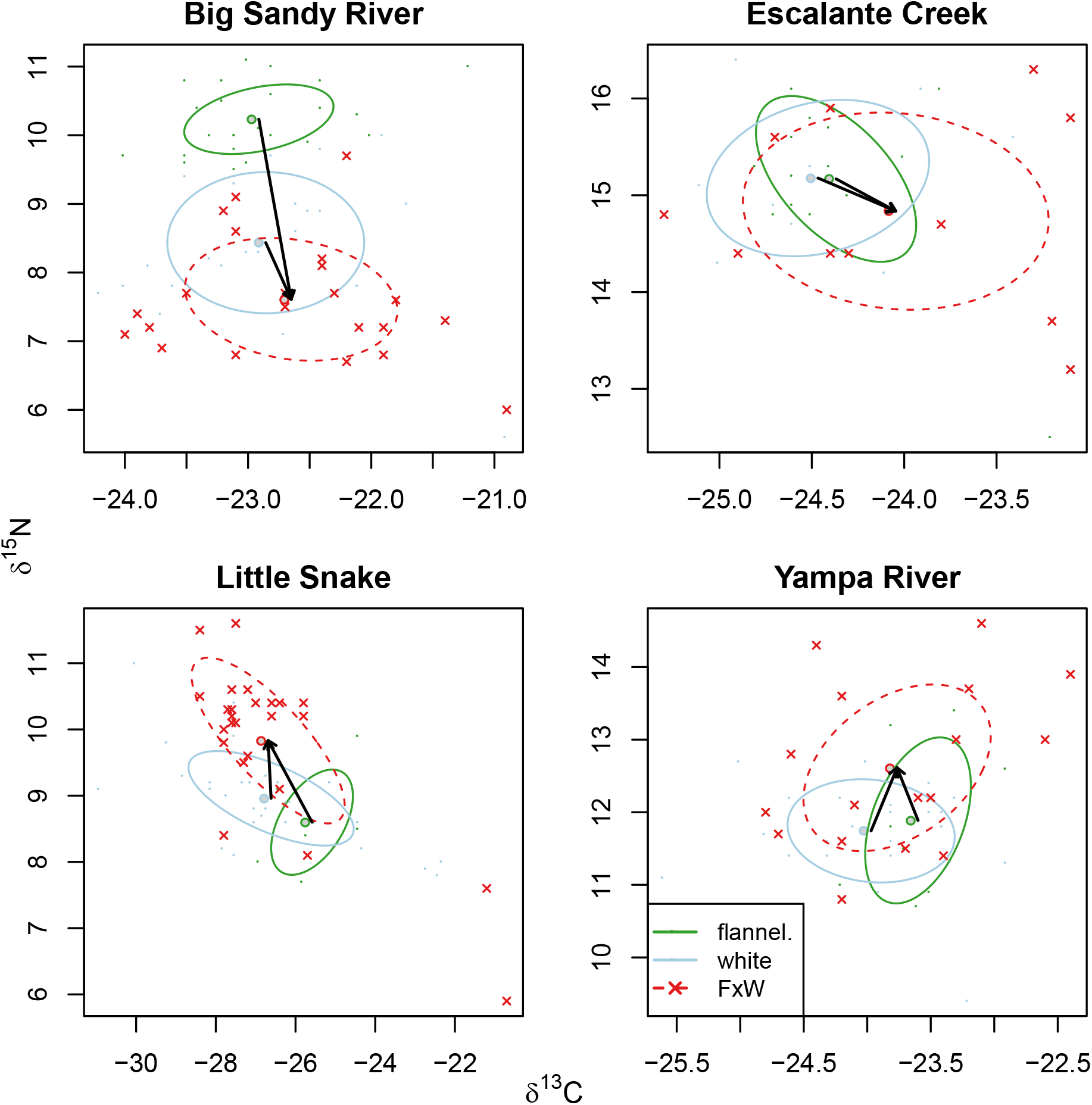
Mean carbon and nitrogen isotopic ratios for flannelmouth and white suckers were similar in most locations. However, as shown by arrows from the mean for parental species to the mean for hybrid individuals, mean values for flannelmouth × white hybrids were transgressive relative to means of parental species. Means are shown with gray points; individual fish are denoted by points color-coded by species or hybrid cross.

Body condition of suckers varied among rivers but not in any particularly predictive way (Fig. 7, Fig. 8). Using Fulton’s condition factor, we compared all species and hybrids. In some locations, white suckers and hybrids had higher body condition than native bluehead and flannelmouth suckers. Relative body condition of species was different in different locations; in some locations (Escalante Creek, White River) bluehead and flannelmouth suckers had similar body condition, while in one location (Yampa River) flannelmouth suckers had significantly better body condition than bluehead suckers. Similarly, in one location all hybrid crosses had similar condition factors to native species (Escalante Creek), while in other locations (Yampa River, White River) flannelmouth*×*white sucker hybrids had higher body condition than at least one native species. These results have some caveats, notably that it is difficult to compare Fulton’s condition factor across species with different body shapes. For this reason, we also used relative weight, W_*r*_, to compare body condition in flannelmouth suckers and white suckers, the two species with existing W_*r*_ equations, in three locations where adequate numbers of both species were sampled. Estimates of W_*r*_ (Fig. 8) contradicted findings using Fulton’s condition factor, and showed that flannelmouth suckers were in significantly better condition than white suckers in one location (Escalante Creek), but in similar condition in the other two rivers where this both species were present. W_*r*_ was similar for flannelmouth suckers and white suckers in the Little Snake River and Yampa River. Taken together, the two metrics of body condition suggest similar ecological success across ancestry classes.

**Figure 7:**
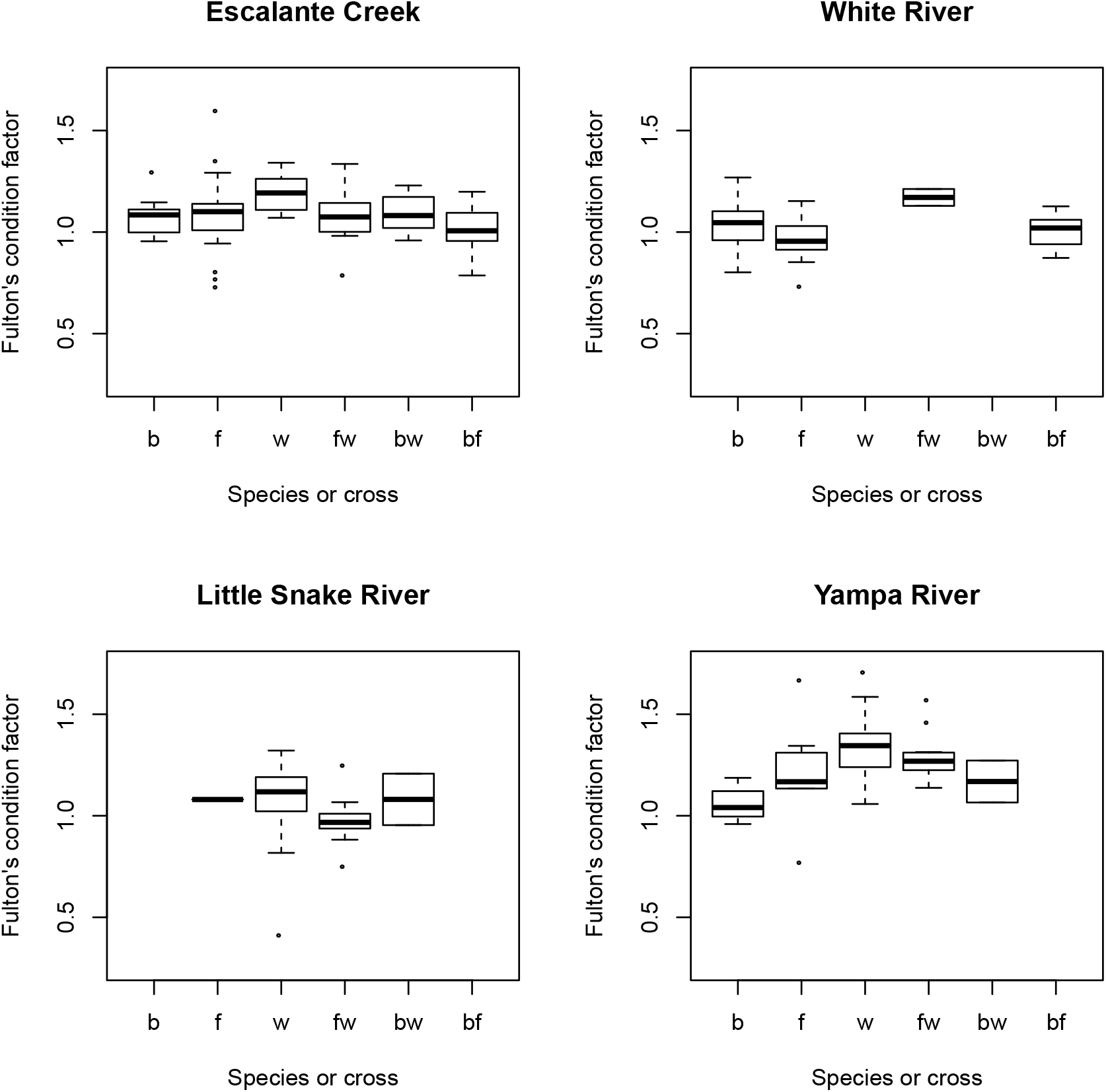
Hybrids and non-native white suckers had similar or higher body condition indices compared to native bluehead and flannelmouth suckers using Fulton’s condition factor. Relative condition of species and hybrid crosses varied across rivers.

**Figure 8:**
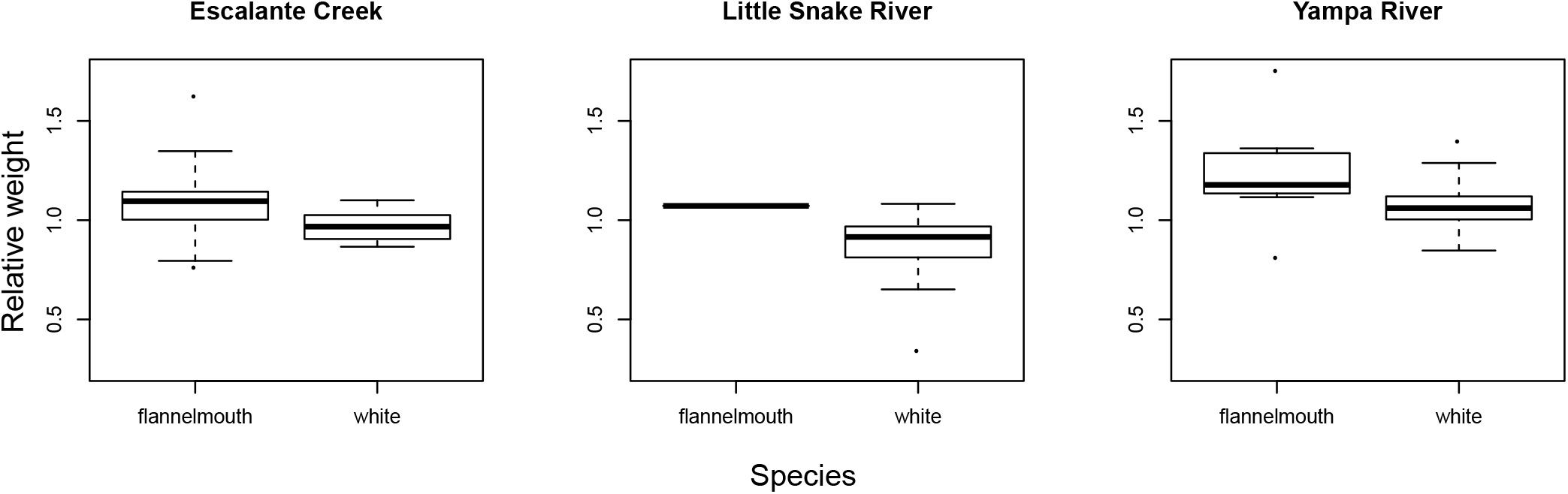
Flannelmouth suckers and white suckers had similar relative weights according to W_*r*_ equations available for these two species. In one river, Escalante Creek, flannelmouth suckers were in better condition according to W_*r*_ estimates; in the other 3 rivers no differences between species were identified.

## Discussion

Ecological relationships between parental species and hybrid crosses varied extensively across the seven rivers in this study. Hybrids interacted differently with the parental species and potentially filled different ecological niches in different locations. The degree of variation across these streams–physical, ecological, and variation in the set of fish taxa–and the relatively low number of locations (seven) made it difficult to identify specific factors associated with variation across streams. However, inclusion of seven locations is a central feature of this study–had we sampled a single location, we would have identified simpler, but not generally applicable, ecological outcomes of hybridization. We did identify an association between ecological crowding and more extensive hybridization between species. It is unclear whether this intensified ecological overlap is a consequence of extensive hybridization, or whether hybridization occurs more extensively in areas with more crowded niche space. Both explanations are plausible and we discuss them in greater detail below.

### Patterns of ecological overlap among species and hybrids

Ecological overlap as measured by isotopic niches (Layman et al. 2007, Newsome et al. 2007) showed high variability among rivers and this variability correlated with the extent of hybridization (Fig. 4). Relationships among species in isotopic space varied across rivers. However, one consistent trend was that bluehead and flannelmouth suckers, the two native parental species, did not overlap substantially in isotopic space (Fig. 2). In three out of the five rivers with greater than one statistically supported cluster, individuals from these two species were largely but not entirely assigned to different clusters in our hierarchical Bayesian clustering algorithm (Fig. 3). We suggest that bluehead and flannelmouth sucker diets generally did not substantially overlap where the species co-occurred. Because both species are historically native to the study locations, it is possible that non-overlapping isotopic niches have evolved to avoid competition. The idea of ecological character displacement in response to competition has a long history in ecology (Connell 1980, Robinson and Wilson 1994), and has previously been invoked to explain evolutionary origins and maintenance of species diversity (Schluter 2000, Seehausen 2007). Our results are consistent with evolution of divergent ecological niches in flannelmouth and bluehead suckers.

In contrast, introduced white suckers overlapped in isotopic niche space with flannelmouth, bluehead, or with both species, suggesting some degree of shared resource use between introduced white sucker and one or both native species at every location sampled. Overlap of white suckers with bluehead suckers in some locations was unexpected because we anticipated white suckers would be more similar ecologically to flannelmouth than to bluehead suckers, due to similar mouth morphology and because white suckers preferentially eat invertebrates (Baxter et al. 1995, Cross et al. 2013, Walsworth et al. 2013). This overlap in isotopic space suggests that different native species might face competition from introduced white suckers in different rivers. Variable position of white suckers relative to the other species in this study also suggests that the resource use of white suckers might be flexible or opportunistic (as in Corse et al. 2009). If this pattern were true, then ecological flexibility and generalist traits of white suckers might have facilitated successful invasion of the Upper Colorado River basin, and might explain to some extent the success of hybrid individuals with white sucker ancestry.

Hybrid individuals also occupied different isotopic space relative to parental species in different locations (Fig. 2). Hybridization of bluehead suckers with both flannelmouth and white suckers produced some hybrids that were intermediate in isotopic ratios relative to parental species (Fig. 5). However, some individual bluehead*×*white and bluehead*×*flannelmouth sucker hybrids also matched parental phenotypes in isotopic space, suggesting similarity of resource use to parental species. Intermediacy in diet might be linked to intermediacy in mouth morphology. Hybrid individuals with bluehead*×*white and bluehead*×*flannelmouth ancestry have intermediate mouth morphology, including a reduced scraping ridge (Hubbs et al. 1943, Hubbs 1955, Quist et al. 2009). Intermediate morphology in hybrids corresponds to intermediate diets in other fish species. Both stickleback and centrarchid fish hybrids use intermediate food sources when they have intermediate feeding morphology (Arnegard et al. 2014, McGee et al. 2015), with variable ecological success depending on the quantity and quality of food resources available. A parental-like feeding phenotype might be advantageous to hybrid individuals if parental phenotypes are optimal for feeding on available resources (as in Arnegard et al. 2014), but also might lead to competition between the native parental species and their hybrid offspring if preferred resources are limited (as in Seiler and Keeley 2009). If overlap in isotopic space does correspond to consumption of similar food resources in these rivers, competition between hybrids and parental species might occur. Competition might therefore be a serious concern for conservation of native *Catostomus* species in locations where hybridization is ongoing and resources are limited. We currently do not know to what extent fish production in these rivers is resource limited, so we cannot infer whether selective pressures associated with competition are strong, but matching ecological phenotypes suggests that competition is possible.

Hybridization can also produce individuals with phenotypes that differ from either parental species and are not intermediate; indeed, novel phenotypes produced by hybridization have promoted evolutionary diversification in diverse groups of organisms (e.g. Rieseberg et al. 2003*a*, Marques et al. 2019, Meier et al. 2019). We observed transgressive isotopic ratios in a subset of flannelmouth*×*white hybrid individuals (Fig. 6). These individuals exceeded values for any other *Catostomus* species in these rivers, including parental species of the hybrids. In Escalante Creek, *δ*^13^C values for a subset of flannelmouth*×*white hybrids were transgressive. In the Yampa River, *δ*^15^N ratios for a subset of individuals were transgressive. Transgressive values for *δ*^13^C likely correspond to use of different carbon resources, potentially in a different part of the river (Finlay 2001). Transgressive values for *δ*^15^N could also correspond to use of a different portion of the river (e.g., feeding in a deep pool where denitrification is occurring), but nitrogen isotope values are more commonly interpreted as a rough indicator of trophic position (Post 2002).

Individual fish with transgressive *δ*^15^N might be feeding on a higher trophic level by eating more invertebrates than flannelmouth or white suckers, or consuming some other resource like larval fish. It is not possible to identify causes of elevated *δ*^15^N without additional data like direct observation of stomach contents and a detailed survey of the base of the food web, which are goals for future study. Examples of transgressive phenotypes in hybrids are well-documented in other organisms (Rieseberg et al. 1999*a*, Stelkens and Seehausen 2009), and hybrids can use different resources and habitats than parents (Rieseberg et al. 2003*b*, Gompert et al. 2006, Rieseberg et al. 2007). In at least one other group of organisms, cichlid fishes, likelihood of transgressive phenotypes is correlated with evolutionary distance (Stelkens et al. 2009), but in *Catostomus* fishes the transgressive hybrids appear to result from hybridization between closely related and ecologically similar taxa (flannelmouth and white suckers). Transgressive phenotypes are unlikely to be fit when they initially arise, as they do not reflect a long history of adaptation, but some fraction of transgressive phenotypes might confer a novel ability to use certain resources or habitats, and have been implicated as a potential source of evolutionary novelty in diversification (Rieseberg et al. 2003*b*, Stelkens and Seehausen 2009). Regardless of specific drivers of elevated *δ*^13^C and *δ*^15^N, transgressive isotopic ratios suggest that some individual hybrids might use different resources than parental species. In both of these rivers, isotopic ratios of hybrids had more among-individual variation than isotopic ratios of parental species, as shown by the slightly larger standard ellipse areas. Consumption of alternative resources could be advantageous in more crowded ecological niche space, such as locations containing many species and hybrids that overlap in dietary requirements.

### Food web effects of hybridization

In rivers with extensive hybridization, niche space occupied by *Catostomus* species and their hybrids was more compressed (Fig. 2), and occupied by more species and hybrids that potentially overlap in resource use, suggesting that one potential effect of hybridization is ecological crowding, with potential for competition. In locations where two parental species, flannelmouth sucker and white sucker, overlapped more in isotopic space, there was less backcrossing and hybridization was mostly constrained to first generation hybrids (Fig. 4). These lines of evidence suggest that although there might be some filtering of hybridization outcomes by environmental conditions and available niche space, it is perhaps more likely that hybridization in *Catostomus* suckers could reshape food webs. Non-native species alter food web dynamics in freshwater fishe communities, including isotopic niche (Sagouis et al. 2015, Rogosch and Olden 2020), but the role of hybrids is less well documented. One study of *Catostomus* ecology that used stable isotopes, Walsworth et al. (2013), showed that introduction of white suckers and other non-native species into tributaries of the Upper Colorado River basin could increase food chain length and increase niche crowding, consistent with our findings here. However, Walsworth et al. 2013 excluded hybrids, which make up a large fraction of the *Catostomus* fish present in many parts of the Upper Colorado River basin (Gill et al. 2007). In our study, it is notable that although the number of identified distinct clusters in isotopic space was correlated with number of parental species (Fig. 4), the same relationship did not hold for number of clusters and number of ancestry classes, i.e., parental species plus hybrid crosses. This finding suggests that hybrids were not likely to be accessing different resources or occupying available niches, except perhaps in the case of individuals with transgressive phenotypes, but rather were overlapping with ecological resource use of parental species, with the potential for competition. It is plausible that the degree of ecological crowding in a river might correspond to increased selection pressures for foraging success, as hybrids, non-native parental species, and native parental species compete for resources and space. Competition and

It is unclear what fitness consequences these interactions between parental species and hybrids might have. Attempts to use body condition as a measure of ecological success and proxy for fitness produced equivocal results, because different measures of body condition produced opposing patterns (Fig. 7, Fig. 8). These contradictory results illustrate some of the methodological challenges associated with estimating body condition (Peig and Green 2010), and suggest caution is warranted in attaching too much weight to body condition as a fitness proxy. A simpler approach, Fulton’s condition factor (*K*), fails to account for species-level differences in body shape. The *Catostomus* species in this study had similar body shapes, but white suckers have a wider caudal peduncle, while flannelmouth and bluehead have more slender caudal peduncles (Baxter et al. 1995). The other approach we used, calculating relative weight (W_*r*_) from preexisting standard weight equations, does account for differences in body shape, but relies on equations developed from reference individuals, and could not be applied to all species, and especially not for hybrids. One particularly troubling aspect of the W_*r*_ approach for flannelmouth suckers was that the standard weight equation was developed using reference individuals from the Upper Colorado River basin (Didenko et al. 2004), where flannelmouth suckers are endemic, and where white suckers and hybrids have long been present. If there are negative effects of competition, they were incorporated into our conception of what a “standard” flannelmouth sucker should weigh due to the limitations of body condition indexes based on reference populations.

From calculations using Fulton’s condition factor, *Catostomus* hybrids were in similar or slightly better body condition than parental species (Fig. 7). Notably, however, the relationship between flannelmouth suckers and white suckers was reversed using W_*r*_, with flannelmouth suckers in slightly better body condition (Fig. 8), which leaves some uncertainty about the accuracy and meaning of these comparisons. However, hybrids must have been at least moderately ecologically successful, because body condition in hybrids was similar to parental species using both metrics. One caveat is that we only sampled adults, which by definition must have been ecologically successful enough to survive to maturity. It is possible that hybrid individuals with less ecologically successful phenotypes were produced, but did not survive early life stages. In some known examples, a broad range of interspecific hybrids can be viable under favorable conditions, but only a subset of hybrid individuals survive in a given environment (as in *Populus* trees; Lindtke et al. 2014). It is possible that a similar filtering of a subset of hybrid genotypes occurred with *Catostomus* hybrids, and that the hybrids sampled as adults were those that survived strong selective pressures on ecological traits as juveniles.

More empirical studies of fitness in hybrid individuals, including *Catostomus* hybrids, will be needed to develop a more mechanistic understanding of hybridization. Clarifying this point is essential because relative fitness of parental species and hybrids has a profound effect on how hybridization affects the evolution of species (reviewed in Arnold M and Hodges 1995, Arnold et al. 2012). In some instances of interspecific hybridization, as in tiger salamanders, hybrids survived at higher rates than either parental species (Fitzpatrick and Shaffer 2007). However, it is important to distinguish between the well-being of individuals—i.e., survival, growth, and ecological success—and reproductive success. Survival and growth cannot be equated with fitness if hybrids cannot reproduce, and there are many examples of hybrid sterility (e.g. Sweigart et al. 2006, Good et al. 2008). There might also be fitness differences between first-generation hybrids and later generation hybrids, particularly if there is heterosis in *F*_1_ hybrids. For example, in native westslope cutthroat trout (*Oncorhynchus clarkii lewisi*) and introduced rainbow trout (*Oncorhynchus mykiss*) hybridization, *F*_1_ hybrids had relatively high fitness, including high reproductive success, but later-generation hybrids had markedly lower fitness (Muhlfeld et al. 2009). Studies of fitness in wild, free-living populations are logistically challenging, but would add to our understanding of the causes and consequences of interspecific hybridization.

### Conservation implications

The common native parental species in this study, bluehead and flannelmouth suckers, are currently the target of extensive management and conservation efforts by state agencies (Gel-wicks et al. 2009, Senecal et al. 2010). Previous work on genomic outcomes of hybridization suggested that hybridization might act as a demographic sink due to reduced conspecific reproduction and apparent absence of hybrids beyond the *F*_1_ generation in some crosses (Mandeville et al. 2015, 2017). We can also now infer some potential ecological effects of hybridization. Hybrids and non-native parental species overlapped substantially in isotopic space with native parental species, which suggests that hybrids and introduced species occupied similar ecological niches to native species, and therefore might compete for resources. Additionally, in some rivers hybrids and non-native species had better body condition than native species by one metric, suggesting that hybrids and introduced species might sometimes exploit resources better than parental species. When combined, these lines of evidence suggest additional potential obstacles to conservation of native species: hybrids and introduced species are likely to compete with native species, but hybrids do not contribute substantially to the gene pool in some rivers. Thus, hybridization might sometimes be both a demographic sink and a resource sink. However, the variability uncovered by both this study and genetic studies of hybridization suggest that these dynamics might vary substantially by river, making outcomes of hybridization more complicated to predict. If we had studied these ecological dynamics in a single river, our conclusions would have been incorrect; it is critical to recognize that ecological outcomes of hybridization, like genomic outcomes, may vary substantially and might require different conservation strategies in different locations.

## Conclusions

In this study, we showed that ecological niches occupied by hybrid individuals varied within and across rivers. Across replicate locations where hybridization occurs, hybrids of similar ancestry had variable feeding ecology, either matching parental species isotopic ratios, intermediate between parental species, or displaying a transgressive isotopic ratio beyond the range of either parental species. This variation might have arisen due to greater plasticity in hybrid feeding phenotypes, or might suggest that postzygotic selection constrained hybrids to the range of phenotypes that would be ecologically successful in a given location. We also suggest that overall food web structure might either influence hybridization, or be altered by the presence of hybrids and non-native species. Together, these results make a compelling case for more study of how ecological interactions can constrain or facilitate hybridization, and how hybridization might alter ecological function of communities and modify fitness landscapes for hybridizing species.

## Acknowledgments

Isotope analysis was funded by the Biodiversity Institute at the University of Wyoming (Biodiversity Research Grant to EGM). Genomic data were generated with funding through the Wyoming Game and Fish Department (State Wildlife Grant #001793 and Bureau of Land Management Cooperative Agreement 12AC20048) and Colorado Parks and Wildlife (Species Conservation Trust Fund project SCTF001C). EGM was supported in part by NIH INBRE funding to the University of Wyoming (NCRR P20RR016474/NIGMS P20GM103432) and by the UW Biodiversity Institute. Computing was accomplished with an allocation from from the University of Wyoming’s Advanced Research Computing Center, on its Mount Moran IBM System X cluster (http://n2t.net/ark:/85786/m4159c) and Teton Intel x86 64 cluster (https://doi.org/10.15786/M2FY47). We also thank the Wyoming Game and Fish Department and Colorado Parks and Wildlife for providing samples and supporting our work on *Catostomus* hybridization, most notably Mark Smith, Kevin Gelwicks, Bobby Compton, and Kevin Thompson. We also thank the staff of the Stable Isotope Facility at the University of Wyoming, especially Chandelle MacDonald, for their assistance and expert advice. This manuscript was improved by comments from the Walters lab at the University of Wyoming and the Mandeville lab at the University of Guelph.

